# Global Landscape of Native Protein Complexes in *Synechocystis* sp. PCC 6803

**DOI:** 10.1101/2020.03.07.980128

**Authors:** Chen Xu, Bing Wang, Lin Yang, Lucas Zhongming Hu, Lanxing Yi, Yaxuan Wang, Shenglan Chen, Andrew Emili, Cuihong Wan

## Abstract

*Synechocystis* sp. PCC 6803 (hereafter: *Synechocystis*) is a model organism for studying photosynthesis, energy metabolism, and environmental stress. Though known as the first fully sequenced phototrophic organism, *Synechocystis* still has almost half of its proteome without functional annotations. In this study, we obtained 291 protein complexes, including 24,092 protein-protein interactions (PPIs) among 2062 proteins by using co–fractionation and LC/MS/MS. The additional level of PPIs information not only revealed the roles of photosynthesis in metabolism, cell motility, DNA repair, cell division, and other physiological processes, but also showed how protein functions vary from bacteria to higher plants due to the changed interaction partner. It also allows us to uncover functions of hypothetical proteins, such as Sll0445, Sll0446, Sll0447 participating in photosynthesis and cell motility, and Sll1334 regulating the expression of fatty acid. Here we presented the most extensive protein interaction data in *Synechocystis* so far, which might provide critical insights into the fundamental molecular mechanism in Cyanobacterium.

## Introduction

Cyanobacteria represent the phylogenetic ancestors of chloroplasts from present–day plants [1,2]. The oxygen generated by oxygenic photosynthesis is believed to change the atmospheric composition and promoting biodiversity on earth [3]. *Synechocystis* is a unicellular photoautotrophic cyanobacterium, which is an ideal model organism for studying photosynthesis, energy metabolism, and environmental stress [4,5]. The genome of *Synechocystis* was sequenced in 1996 [6], and its proteome has also been well analyzed in the last two decades [7,8]. However, about two-thirds of its proteome in UniProt are “hypothetical protein”, and most of which lack functional annotation.

Completions of many important biological functions rely on stable physical interactions between two or more proteins. Protein interaction is critical to understanding the fundamental molecular biology of organisms, which can be used for predicting annotation, finding new drug targets, etc. [9]. However, information on *Synechocystis”*s PPIs is quite a deficiency. Researchers have tried to analyze PPIs by using yeast two–hybrid (Y2H) assay and several kinds of prediction algorithms [10–13]. However, in the STRING database, only 6510 PPIs of *Synechocystis* involved in 1876 proteins were annotated with “experiments” until January 2019, which contained PPIs that relevant information transferred from other organisms. Even for those well– known *Synechocystis* protein complexes, for example, Photosystem II (PSII) complexes that still need to identify novel assembly factors to understand its biogenesis better [14,15]. The phototactic movement of cells was influenced by motility apparatus and light, but the link protein between photoreceptors and the motility apparatus remains uncertain [16,17]. Thus, it is demanding to globally identify PPIs of *Synechocystis*, providing a useful resource to the biology community.

The high–throughput methods have applied to systematically determine global protein interaction maps in many model organisms, such as *E.coli*, fly, worm, yeast, and human [18–21]. Several techniques have been developed to identify protein complexes at proteome scale, e.g., Y2H, affinity purification mass spectrometry (APMS), and co–fractionation coupled with mass spectrometry (CoFrac–MS). Among these methods, CoFrac–MS can rapidly detect hundreds to thousands of stable protein complexes involved in interactions of multiple proteins under native conditions [22]. CoFrac-MS assay has been broadly used to globally identify PPIs [23–25].

To reveal more protein interactions and uncover a clue of their biological functions, we applied CoFrac-MS to analyze the protein complexes of *Synechocystis*. In this work, we predicted 291 protein complexes containing 24,092 highly confident PPIs among 2062 proteins. This network facilitates our comprehensive understanding of the relationship between photosynthesis and other functions, such as carbohydrate metabolic process, signal transduction, ion transport, cell division, transcription, etc. In addition, we applied the protein interaction information to predict and confirm the new functions of proteins, such as Sll0445, Sll0446, Sll0447, and Sll1334. This work allows us to comprehensively understand the fundamental molecular organization and mechanism of *Synechocystis* and other Cyanobacteria species.

## Results and discussion

### Workflow for protein complexes identification in *Synechocystis*

The experimental workflow is similar to previous work **(Figure 1A)** [20]. Protein mixtures were separated and analyzed by ion-exchange chromatography (IEC), size–exclusion chromatography (SEC), or sucrose density gradient centrifugation (Suc–DGC). In total, 181 fractions were collected, and 2906 proteins were identified (**Table S1**). Proteins in lysis samples were separated effectively according to their molecular weight (MW) or pI **(Figure S1)**. After fractionation, samples were analyzed using LC/MS/MS. The value of Rapp, the ratio of Mapp (proteins apparent molecular mass, **Figure S2**) to Mmono (the predicted monomeric mass), was evaluated to estimate whether a protein involved a stable complex on SEC column [26,27]. Rapp ≥ 2 is used to classify proteins to be not oligomeric but within a complex, while Rapp ≤ 0.5 suggests protein degradation during the protein extraction process (**Figure 1B**).

**Figure 1.**
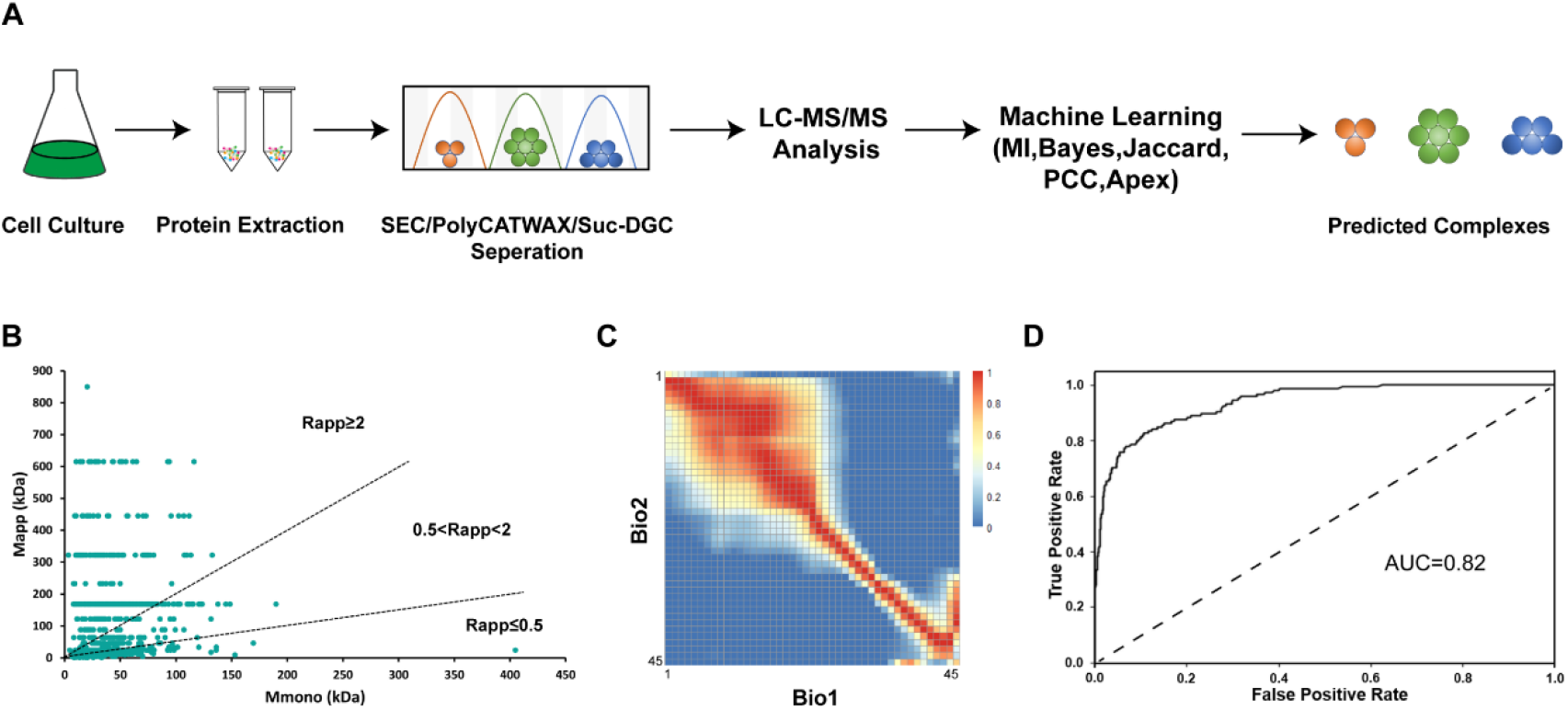
Workflow for the identification of native protein complexes. **A.** Schematic diagram of co–fractionation LC/MS/MS and machine learning. Lysates were produced containing a mixture of protein complexes, which were separated by SEC, IEC or Suc–DGC. Proteins in each fraction were digested with trypsin and analyzed by nano–LC–MS/MS. Putative PPIs were predicted by machine learning using EPIC toolkits. **B.** The calculation of Rapp, which can accurately reflect the oligomerization state of proteins. Rapp, the ratio of Mapp to Mmono; Mapp, apparent molecular mass; Mmono, predicted molecular mass of monomer. A protein with Rapp ≥ 2 means it has interaction with other proteins. **C.** Heatmap of the Pearson correlation coefficients of the protein quantification signals in two SEC biological replicates. **D.** Receiver operating characteristic curve of machine learning.

The SEC is advantageous to provide direct distribution of protein in different MW [24]. However, SEC is not sufficient to separate the protein complexes with MW beyond its valid separation range. To improve protein separation efficiency, we applied IEC as an additional separation technique. These two techniques are complementary in the way that IEC can separate protein complexes that are not distinguishable in SEC **(Figure S3)**. The reproducibility between two biological replicates was confirmed by Pearson correlation coefficients using spectral counts of identified proteins **(Figure 1C)**.

The stable protein complexes bind tightly and can be detected by CoFrac–MS, while the unstable protein complexes might present unsatisfactory correlation profiling [19]. For example, photosystem components, NAD(P)H–quinone oxidoreductase, RubisCO complexes, and C–phycocyanin, tend to have a consistent correlation profiling **(Figure S4)**. These protein elution profiles in turn confirm that CoFrac–MS is a powerful tool to explore global protein interactions in organisms. We confirm the final set of PPIs by machine learning using Elution Profile–based Inference of Complexes (EPIC) [28], with collected 47 gold standard complexes from UniProt and IntAct database **(Table S2)**. The output of machine learning was well organized and ready for subsequent analysis to predict protein complexes **(Figure 1D; Figure S5)**.

### Photosynthetic apparatus involved in multiple metabolic pathways

In our datasets, 2214 proteins participate in 35,028 highly confident protein pairs that were produced by machine learning **(Table S3)**. About 10% of PPIs have an overlap with the published database, including STRING and IntAct **(Figure 2A)**. Our study is the most extensive PPIs data in *Synechocystis* so far, which can provide evidence for further functional study in the future.

**Figure 2.**
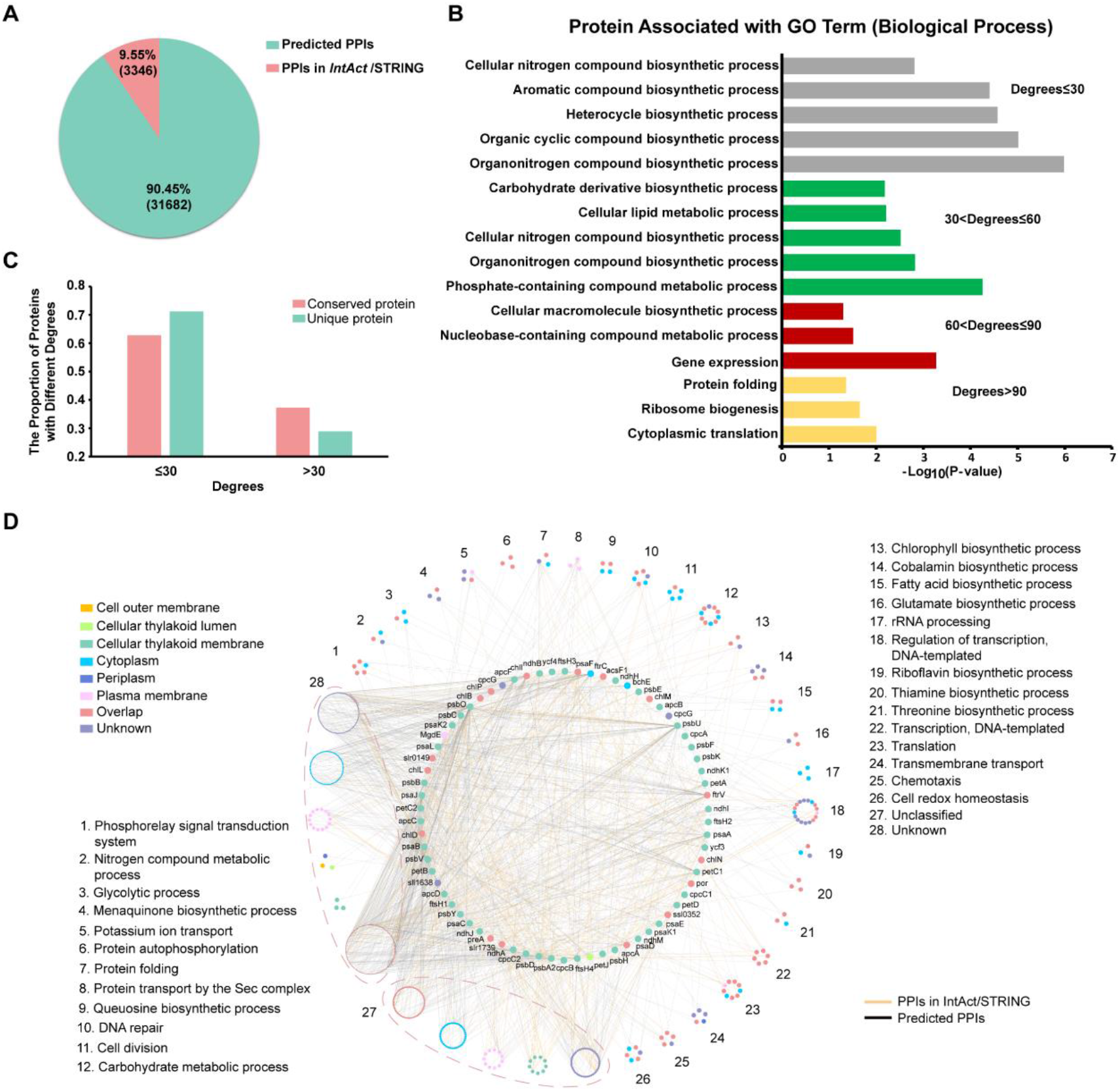
The characteristic of predicted PPIs. **A.** Pie chart showing the overlaps of predicted co–complex PPIs with PPIs from IntAct and STRING databases. **B.** Gene Ontology analysis of proteins with different degrees, which is the number of edges that one protein links to other proteins in the network. (P < 0.05) **C.** The degree distributions of conserved proteins and unique proteins in *Synechocystis*. The conserved protein means it has homologous with protein in *E.coli, A.thaliana, H.sapiens*, and *S.cerevisiae*. **D.** The photosynthesis interacted proteins were classified by function. Each section has three or more proteins aggregation with the same GO term. Different node colors represent the subcellular localization of the protein.

Degree, the number of proteins that participated in the interactions, can reflect the critical role of this protein in the network [29]. We compared the distribution of protein degrees in *Synechocystis* with several different model organisms, such as *E.coli, S.cerevisiae, A.thaliana*, and *H.sapiens*, according to the Mentha database [30]. We observed that most of the proteins tend to have low degrees, and these proteins tend to be annotated with different metabolism pathways in *Synechocystis*, for example, organonitrogen, aromatic, and cellular nitrogen compound biosynthetic process **(Figure 2B; Figure S6)**. In contrast, the ribosomes and heat shock proteins present high degrees in our dataset and are conservative across different species (Figure 2B). The ribosomes and heat shock proteins, such as DnaK1 and DnaJ, are usually highly conserved during species evolution and occupy an important position in metabolism [31]. The conserved proteins tend to have high degrees [32]. Additionally, the proteins that have homologous in *E.coli, A.thaliana, H.sapiens*, or *S.cerevisiae* were found to have higher degrees than the proteins that unique in *Synechocystis* **(Figure 2C)**.

As a model organism to study photosynthesis, *Synechocystis* has a classical photosystem structure containing Photosystem I (PSI), PSII, Cytochrome b_6_f complexes, and ATP synthase complexes [5]. There are 70 proteins annotated with photosynthesis and 760 proteins were found to have direct interactions with them in our dataset **(Table S4)**. Photosynthesis associated proteins were clustered into 26 different groups based on their functions, including phosphorelay signal transduction system, potassium ion transport, DNA repair, cell division, carbohydrate metabolic process, transcription, DNA–templated, translation, chemotaxis, and cell redox homeostasis **(Figure 2D)**. The results indicate how photosynthesis influences many critical biological processes other than photosynthetic carbon fixation. For example, we observed that several CheA–like proteins interact with photosynthetic core proteins in our database. The CheA-like proteins contain phosphorelay sensor kinase activities and participate in chemotaxis in cells [16]. It suggested that cells might control motility direction by regulating the state of CheA-like proteins through phosphorelay in photosynthesis. The protein transport by Sec complexes also had an association with photosynthesis proteins. The proteins SecD and SecF might participate in the PSII assembly process by interacting with different PSII core proteins with a similar function as SecY protein (Figure 2D). SecY is the main transmembrane subunit of preprotein translocase that is essential for PSII assembly by interacting with YidC insertase to facilitate co-translational pD1 insertion [33,34]. In our database, the protein ChlD was also observed to interact with SecD, which indicated that it might be functional to deliver chlorophyll to the newly synthesized pD1, carrying a similar role as protein ChlG (Figure 2D) [34]. Besides, we observed that proteins interacting with photosynthetic proteins are mainly localized in the cytoplasm, while this class of proteins has possible mobility or secretion according to localization identification results [35]. The results provided a clue for understanding how photosynthetic proteins to transmit signals and influence the physiological metabolism in living cells.

### The landscape of native protein complexes in *Synechocystis*

With the highly confident protein pairs assignment from machine learning, we predicted 291 protein complexes with 24,092 PPIs by using the ClusterONE algorithm [36], which is implemented in EPIC software. The set of predicted putative complexes contains well known and highly–conserved complexes such as PSI protein complexes, RubisCO complexes, and NADH dehydrogenase **(Figure 3; Table S5)**. The proteins were shown in different colors according to their subcellular location, while most of the complex components have multiple annotated locations, and marked as “Overlap”. The functions of predicted complexes are involved in photosynthesis, carbon absorption, nitrogen fixing, and electron transfer. Besides, we also observed some proteins with unclear molecular function annotations formed stable protein complexes, such as CRISPR3 system proteins Sll7085–Sll7090, photosystem proteins Sll0144–Sll0149, and ATP–dependent zinc metalloprotease FtsH complexes (Figure 3).

**Figure 3.**
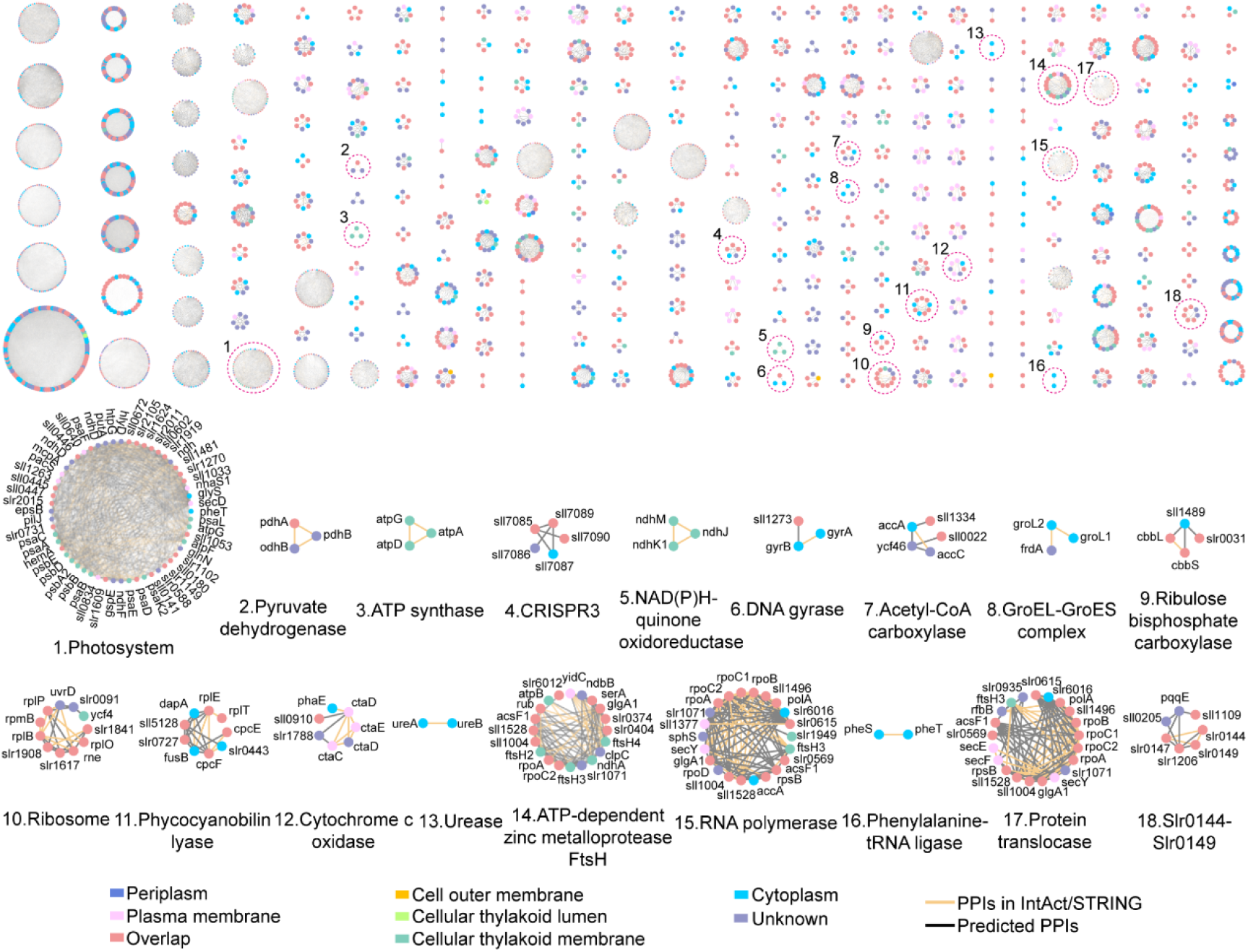
Protein complex map of *Synechocystis*. The top part shows the global landscape of inferred 291 *Synechocystis* protein complexes. The protein nodes are colored according to their subcellular locations. The bottom part presents some of the known protein complex clusters, such as photosystem, ATP synthase, Ribosome, which are annotated with their name or abbreviation. Yellow lines between proteins indicate interactions found in public databases, while gray lines demonstrate the interactions are reported only in this work.

According to computational analysis, *Synechocystis* has three types of CRISPR– Cas system, but their exact functions remain unclear [37–39]. In our dataset, we found that some proteins of CRISPR3 can form stable complexes, and the protein interactions between different CRISPR systems were also observed **(Figure S7)**. These results suggest that different types of CRISPR systems can coordinate with each other to defend cells from virus and plasmid rather than perform function independently. The hypothetical proteins Slr0144–Slr0149 are located on the thylakoid membrane and contain putative bilin binding domains, 4–vinyl reductase (V4R) domains, and 2Fe–2S cluster binding domains. They are mainly involved in the function of photosynthetic repair [40]. The Slr0144–Slr0149 protein interactions are validated by APMS **(Figure S8)**. Among the predicted complexes, we found that the hypothetical protein, Sll0445, Sll0446, and Sll0447 formed protein complexes with pilus assembly proteins, Slr2015 and Slr2018 and photosystem complexes. The physical interactions between Sll0445 and photosynthetic proteins verified by the APMS experiment. The APMS also obtains other partners that inconsistent with CoFrac-MS **(Table S6)**. That might because different methods tend to catch different portions of physical interactions [41]. It is worth mentioning that the proteins Slr2015 and Slr2018 can interact with Sll0445 (Figure 3). The Slr2018 is located at the plasma membrane and regulated by SYCRP1, which is a cAMP receptor protein and influencing the cell motility in *Synechocystis* [42]. The protein Slr2018 shows no homology with other known proteins, but its gene is adjacent to genes *slr2015, slr2016*, and *slr2017*. These genes have roles in pilus morphology and motility, and their proteins were predicted to form protein complexes with photosynthetic proteins (Figure 3) [43]. The results reveal a close connection presents between photosynthesis and cell motility in *Synechocystis*.

### Conservative analysis of *Synechocystis* protein complexes

Half of our predicted protein complexes are conserved with homologous in other species, like *A.thaliana* or *E.coli*, while the rest of them are unique in *Synechocystis* **(Figure S9)**. The predicted protein complexes in *Synechocystis* have a higher similarity with *A.thaliana* and *E.coli* compared to *H.sapiens* and *S.cerevisiae*. Because *Synechocystis* has a close relationship with *A.thaliana* and *E.coli* for the metabolic feature of Gram–negative bacteria and photosynthesis of green plants. The conservativeness of protein complexes was evaluated by the number of proteins with known homologies in each complex. There are 920 proteins and 813 proteins with homologous in *E. coli* and *A.thaliana*, respectively **(Table S7)**. To estimate the evolutionary relationship of *Synechocystis* with *A. thaliana* and *E.coli*, we separated our predicted complexes into different parts according to their homologous proteins. The components of the predicted complexes were divided into four parts, homologous in *E.coli*, homologous in *A.thaliana*, homologous in both *E.coli* and *A.thaliana*, and unique in *Synechocystis* **(Figure 4A)**. Proteins having homologous in *E.coli* and *A.thaliana* retain functions as energy metabolism, organic synthesis, protein expression, and regulation, which are considered as basic activities to be conserved across most species. Proteins with these functions play important roles in maintaining the necessary life activities. Proteins with homologous in *A.thaliana* are mainly related to photosynthesis and pigment biosynthetic process, which is consistent with the fact that one of the most remarkable features shared between *Synechocystis* and *A.thaliana* was photosynthesis. Proteins with homologous in *E.coli* were primarily associated with the polysaccharide metabolic process, protein transport, and coenzyme biosynthetic process, since both *Synechocystis* and *E.coli* belong to Gram - negative bacteria. Cyanobacteria and bacteria share a high sequence homology as evidenced by 16S rDNA analysis. Thus, they have a large number of similarities in cellular structure and physiological properties [44,45].

**Figure 4.**
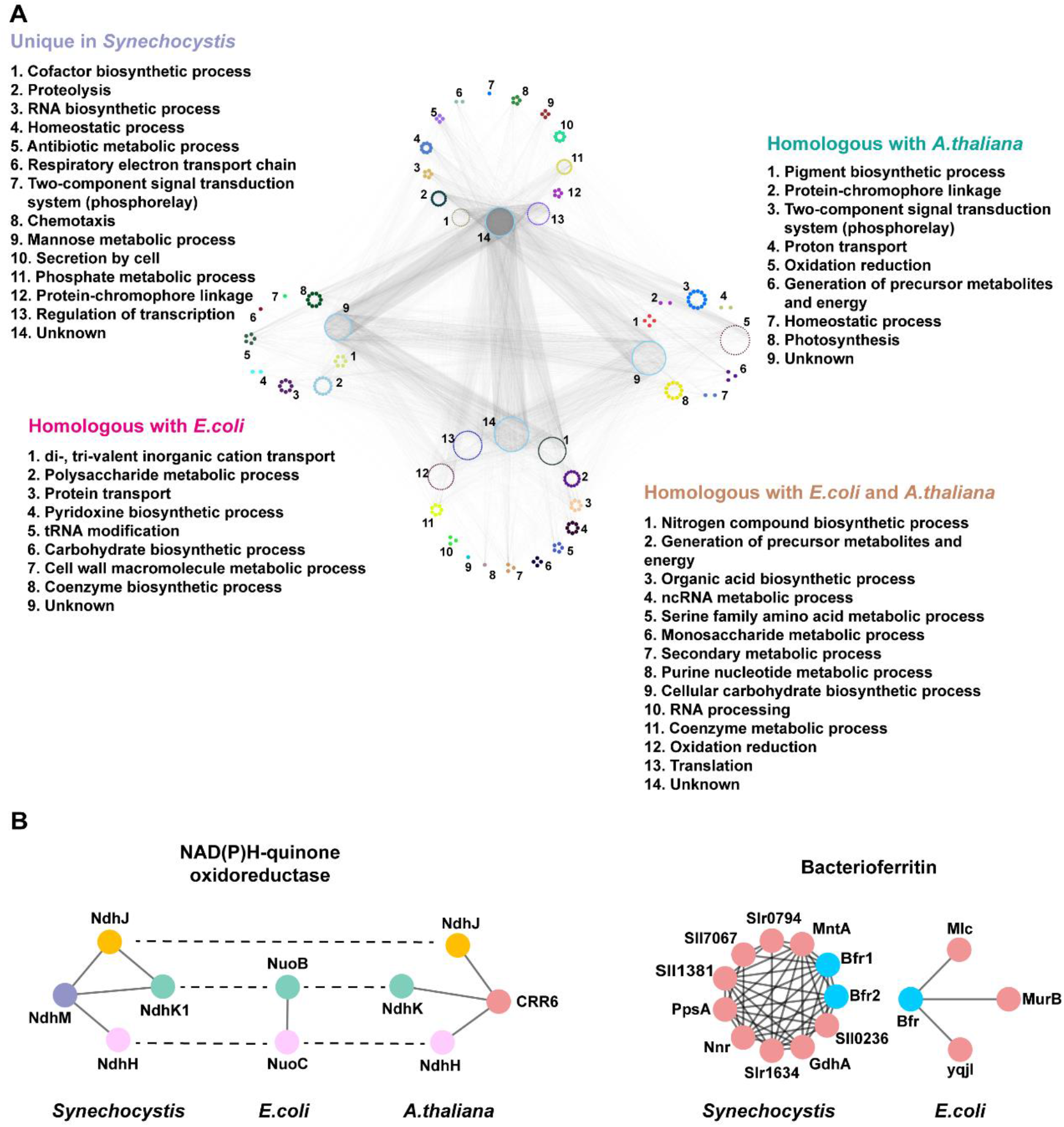
Evolutionarily conserved cyanobacterial protein complexes. **A.** Proteins that have homologous with either *E.coli* or *A.thaliana*, homologous with both *E.coli* and *A.thaliana*, or unique in *Synechocystis* were grouped according to their representative biological processes. **B.** Example of NAD(P)H–quinone oxidoreductase and bacterioferritin protein–protein interaction variation across species. Physical interactions in *E.coli* and *A.thaliana* were collected from the Mentha database. InParanoid orthologs of all three species are depicted with the same colors. Those proteins without homologs in other species are shown in red.

It is noteworthy that the function oxidation–reduction, homeostatic process and two-component signal transduction system (phosphorelay) can be elucidated in more than one of these four groups. However, the proteins that performed these functions and their interaction partners were different (Figure 4A). For example, the proteins annotated as “oxidation-reduction” have homologous in *A.thaliana* and *E.coli* mainly participate in the “generation of precursor metabolites and energy” and cellular respiration processes. While the function of proteins annotated as “oxidation-reduction” with homologous in *A.thaliana* has changed and is mainly associated with photosynthesis **(Figure S10)**.

The localization of NAD(P)H-quinone oxidoreductases (NQOs), which involves electron transfer and shuttles electrons from electron donors to quinones, is different in *E.coli*, *A.thaliana*, and *Synechocystis* [46–48]. The NQOs are on the plasma membrane and participate in the respiratory process in *E.coli*, where the same complexes are found on the thylakoid membrane in *A.thaliana* and *Synechocystis*, indicating that they were involved in the process of photosynthetic electron transport [49,50]. Here we identified 14 NQQs protein **(Table S1)**, and 11 of them form complexes in our final protein network **(Table S5)**. However, the components and functions of NQOs are not entirely consistent in different species [51]. We compared partial NQOs protein interactions of *E.coli*, *A.thaliana*, and *Synechocystis*. We found that the protein NdhM is only found in *Synechocystis* and the proteins NdhJ only exists in *A.thaliana* and *Synechocystis*. In contrast, NdhK and NdhH were conserved in three species **(Figure 4B)**. The results indicated that some components of NQOs participate in the respiratory electron transport and other components may evolve new functions. For example, the proteins of NdhJ and NdhM might be involved in photosynthesis in *A.thaliana* and *Synechocystis*. The appearance of oxidation–reduction in parts of homologous with *A.thaliana* can be a result of adapting to photosynthetic damage or participates in electron transport in *Synechocystis*. Several reactive oxygen species (ROS) produced by aerobic metabolism in photoautotroph were disposed of by antioxidants [52,53]. The presence of new proteins with oxidation–reduction function can effectively reduce the influence of oxidative stress.

Bacterioferritin is an iron–storage protein, whose ferroxidase center binds to and oxidizes Fe^2+^ iron to Fe^3+^ by oxygen. The complexes of bacterioferritin were studied and compared across three species. The results show that there is no ortholog in *A.thaliana* and only one ortholog in *E.coli* (Figure 4B). It is suggested that other proteins might perform this function in *A.thaliana*. The photoautotrophic microorganism tends to have a massive demand for inorganic ions, such as Fe and Mn. Because inorganic ions are usually deficient in the open ocean, a unique mechanism for the microbe to transport and store ions is required. [54,55]. The presence of two orthologs of bacterioferritin in *Synechocystis* indicates that *Synechocystis* may have an additional requirement for irons compared to *E.coli* (Figure 4B). The protein components of bacterioferritin complexes in *Synechocystis* have annotated functions of irons transport that are absent in *E.coli* (Figure 4B). Like bacterioferritin and NQOs mentioned above, the conservative analysis of predicted complexes allows us to explore the biological process within cells and how organisms adapt to environments.

### New functions of hypothetical proteins Sll0445-Sll0447 and Sll1334

Figure 3 indicates that hypothetical protein Sll0445, Sll0446, and Sll0447 have interactions with pilus assembly proteins and photosystem complexes. Sll0445 contains a Tubulin_2 domain, a member of clan Tubulin **(Figure S11A)**. This clan Tubulin subunits served as cytoskeletal elements that were vital for cell division and material transport in all eukaryotes [56]. Sll0446 contains a FtsA domain, which is an actin-like ATPase domain and co-localizes to the septal ring with FtsZ **(Figure S11A)** [57]. The Sll0447 contains a DivIC domain, which is necessary for the formation of both vegetative and sporulation septum **(Figure S11A)** [58]. We constructed interruption mutants for these three genes, respectively, with the insertion of a chloramphenicol–resistance cassette (Cm^R^) into their Open Reading Frames (ORFs) **(Figure S11B)**. Then the abilities of phototactic motility were tested in *Δsll0445, Δsll0446, Δsll0447*, and the wild type. Under the unidirectional light condition, the wild type showed obvious movement tendency to the light, while the *Δsll0445, Δsll0446, andΔsll0447* exhibited impaired ability of phototaxis **(Figure 5A)**. The transcriptome analysis demonstrated that the transcriptional level of *sll0447* was decreased under the mutant of Sycrp1 [42], a gene involved in the phototactic movement [59], indicating that the *sll0447* was regulated by the Sycrp1 and influenced the cell motility.

**Figure 5.**
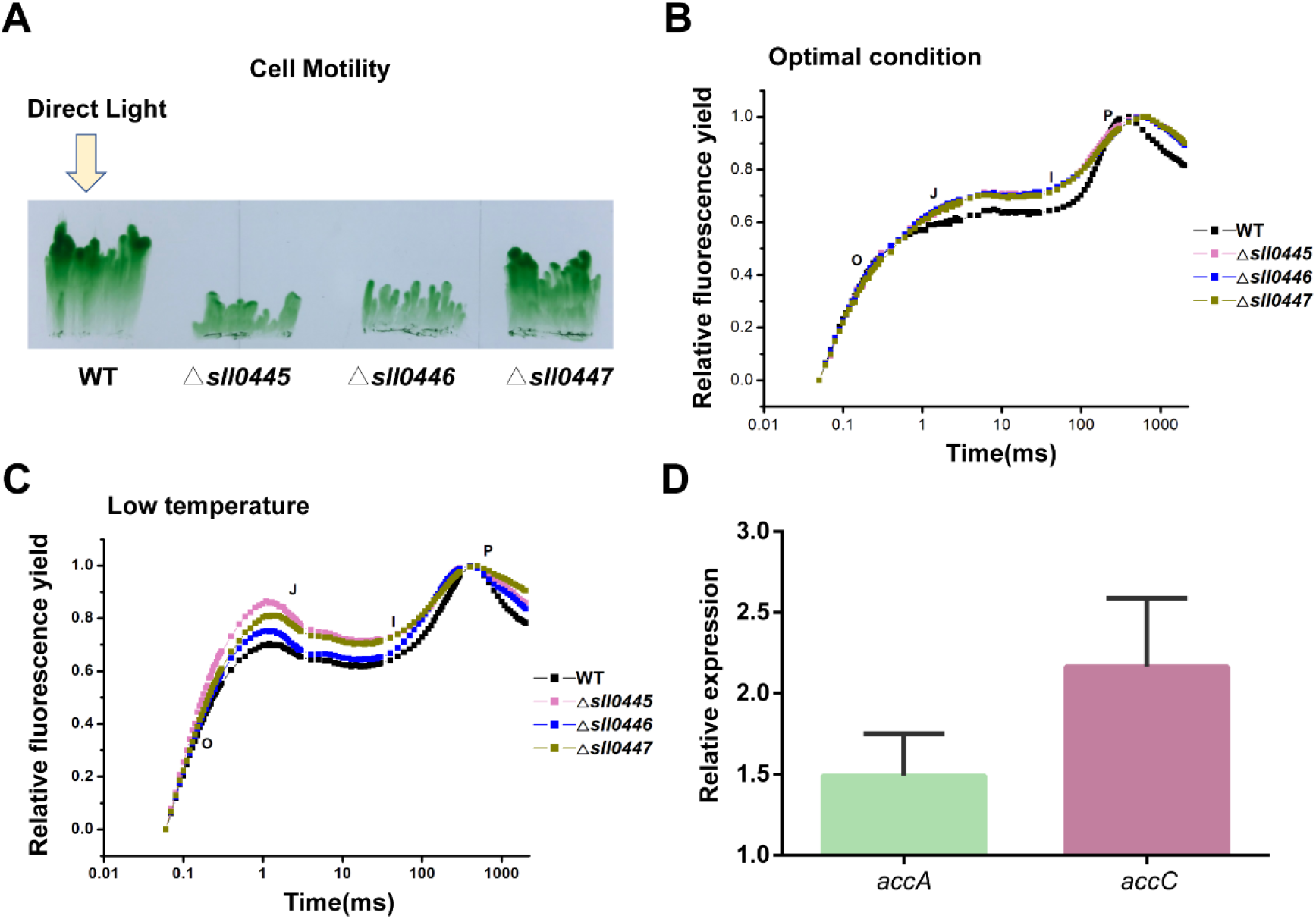
Validation of protein new functions related to cell motility and lipid metabolisms. **A.** The motility state of wild type and mutants under unidirectional illumination (white light). The yellow arrows indicate the direction of the light source. **B-C.** OJIP curve under optimal conditions or low–temperature conditions. The J, I and P steps occurred at about 2 ms, 30 ms, and 400 ms, respectively. **D.** The transcript levels of acetyl coenzyme A carboxylase subunits by qRT–PCR in the wild type strain and *Δsll1334*. Ct–values for each gene were normalized to rnpB.

The hypothetical protein, Sll0445, Sll0446, and Sll0447, not only can form protein complexes with pilus assembly proteins, but also interact with photosystem complexes. The removal of these genes from *Synechocystis”*s genome can affect electron transport of PSII under optimal conditions or low-temperature conditions (20°C) **(Figure 5B and 5C)**. *Synechocystis* exhibits a typical fluorescence induction polyphasic rise called the OJIP curve, which similar to the previous descriptions in plants, green algae, and cyanobacteria [60]. Figure 5B and 5C show sharply increases in the fluorescence yields of mutants at phases J than the wild type, which suggested that the reduction rate of Q_A_ and the oxidation rate of Q_A_-were influenced by genetic deletion of *sll0445, sll0446*, and *sll0447*. The results demonstrate that the proteins encode by *sll0445, sll0446*, and *sll0447* are necessary for optimal function of photosystem and cell motility. However, it is still noteworthy that the pilus assembly proteins distributed on the plasma membrane and photosynthetic proteins localized on the thylakoid membrane. The difference of these proteins in spatial distribution causes the difficulties for proteins that localize on the thylakoid membrane to directly affect a protein complex distributed on the plasma membrane unless a specific protein complex was formed. So far, we know that part of the respiratory electron transport chain in *Synechocystis* located in the thylakoid membrane and partially overlapped with the photosynthetic electron transport chain [49]. In our dataset, the PSI subunits can form a protein complex with NADH dehydrogenase as part of the respiratory chain complexes. The main purpose of *Synechocystis* motility is to obtain plenty of light and consume much energy during chemotaxis. Presumably, the proteins of Sll0445, Sll0446, and Sll0447 may link the structure of the thylakoid membrane with the plasma membrane to adjust the distribution of light on the photosystem and chemotaxis.

A histidine kinase protein Sll1334 interacted with acetyl coenzyme A carboxylase (ACC) **(Figure 3)**. Sll1334 contains GAF domain, which can sense and respond to light signals **(Figure S11A)** [61]. We also observed that the transcript levels of ACC complex were upregulated in *sll1334* interruption mutant compared with wild type strain **(Figure 5D)**. ACC complex catalyzes the acetyl–CoA to form malonyl–CoA, which is part of lipid metabolism and conserved across different species [62]. Okada *et al*. have studied that *sll1334* may function as a suppressive regulator in the cascade and influence cell growth and gene expression involved in glycometabolism under dark conditions [63]. The mechanism is still unclear as to how cyanobacterium adapts to dark conditions and uses glucose as the carbon source for growth. However, our results may provide new insight into this process. Carbohydrate and lipid metabolisms were considered as central metabolism in living cells and had a close relationship with each other to control basic vital activities. Okada *et al*. also have found that *sll1330*, a histidine kinase gene, is located at the upstream of *sll1334* and influences the expression of *sll1334* [64]. The *Δsll1330* and *Δsll1334* did not grow well either under light activated heterotrophic or dark heterotrophic conditions compared to the wild type. However, they could grow as well as the wild type under photoautotrophic conditions [63,64]. These results indicated that carbohydrate and lipid metabolisms in *Synechocystis* were regulated by Sll1330 and Sll1334, and the light was an essential factor to influence this process. Cyanobacteria can produce different bilin–binding photoreceptors when sensing various wavelengths of light, which adjust many essential cellular processes like growth, phototaxis, and photosynthesis to environmental light conditions [61]. The amino– terminal region of photoreceptors protein was found as the photosensory module consisted of a ‘knotted’ structure including a GAF domain, PAS domain, and PHY domain, but not all proteins with GAF domain are photoreceptors. Thus, it needs further confirmation whether Sll1334 is a photoreceptor and how it regulates the ACC complexes expression by sensing light.

Finally, through integrating our data with the cyanobacterial metabolism pathway, we depicted a more comprehensive working model of the phototaxis regulation, PSII assembly and physiological metabolism in *Synechocystis* **(Figure 6)**. Light is one of the essential elements to sustain life, especially for cyanobacteria and green plants that can fix carbon from the outer environment by photosynthesis. In order to obtain enough light, cyanobacteria have been evolved with multiple abilities to adapt to light changes. For example, light energy can be harvested by large antenna complexes– phycobilisomes and phototaxis is capable of helping *Synechocystis* locate an ideal place to collect light [16,65]. The phototaxis can be influenced and regulated by light, the concentration of cAMP, and the structure of pilus in *Synechocystis* [16]. The Slr2015– Slr2018 is one class of proteins involved in the process of pilus assembly and can be induced by cAMP receptor protein Sycrp1 [42]. The hypothetical proteins Sll0445, Sll0446, and Sll0447 are also induced by Sycrp1 and interact with protein pilus assembly proteins, Slr2015 and Slr2018 to regulate cell motilities **(Figure S8)** [42]. Besides, the proteins Sll0445, Sll0446, and Sll0447 can also influence photosynthesis through interactions with photosynthetic core proteins, revealing a close relationship between photosynthesis and cell motility in *Synechocystis* (Figure 5B and 5C). We also provided more detailed information during the PSII assembly processes by combining our predicted PPIs with published data (Figure 6). The biogenesis of PSII required coordinated incorporation of at least 20 protein subunits and a range of organic and inorganic cofactors [14]. Some of the proteins were well understood during pratA–dependent PSII assembly processes. For example, proteins SecY, YidC, and CtpA can facilitate D1 maturation from pD1; proteins ChlG, HliD, and Ycf39 were involved in the process of delivering the chlorophyll to new D1 or pD1 [34,66,67]. In addition to these proteins, we observed the proteins SecD, SecF, and ChlD presented a good elution profiling with these proteins, which indicated that Sec complexes and ChlD might play a critical function in the maturation of PSII during pratA-dependent PSII assembly processes (Figure 6). *Synechocystis* can survive under photoautotrophic or heterotrophic growth conditions. The protein Sll1334 can regulate the expression of genes involved glucolipid metabolism and GAF domain in Sll1334 may have an important role in controlling the utilization of sugars and lipids during heterotrophic growth. These proteins could expand our understanding of the regulation of cell motilities, PSII assembly, and glucolipid metabolism in *Synechocystis*.

**Figure 6.**
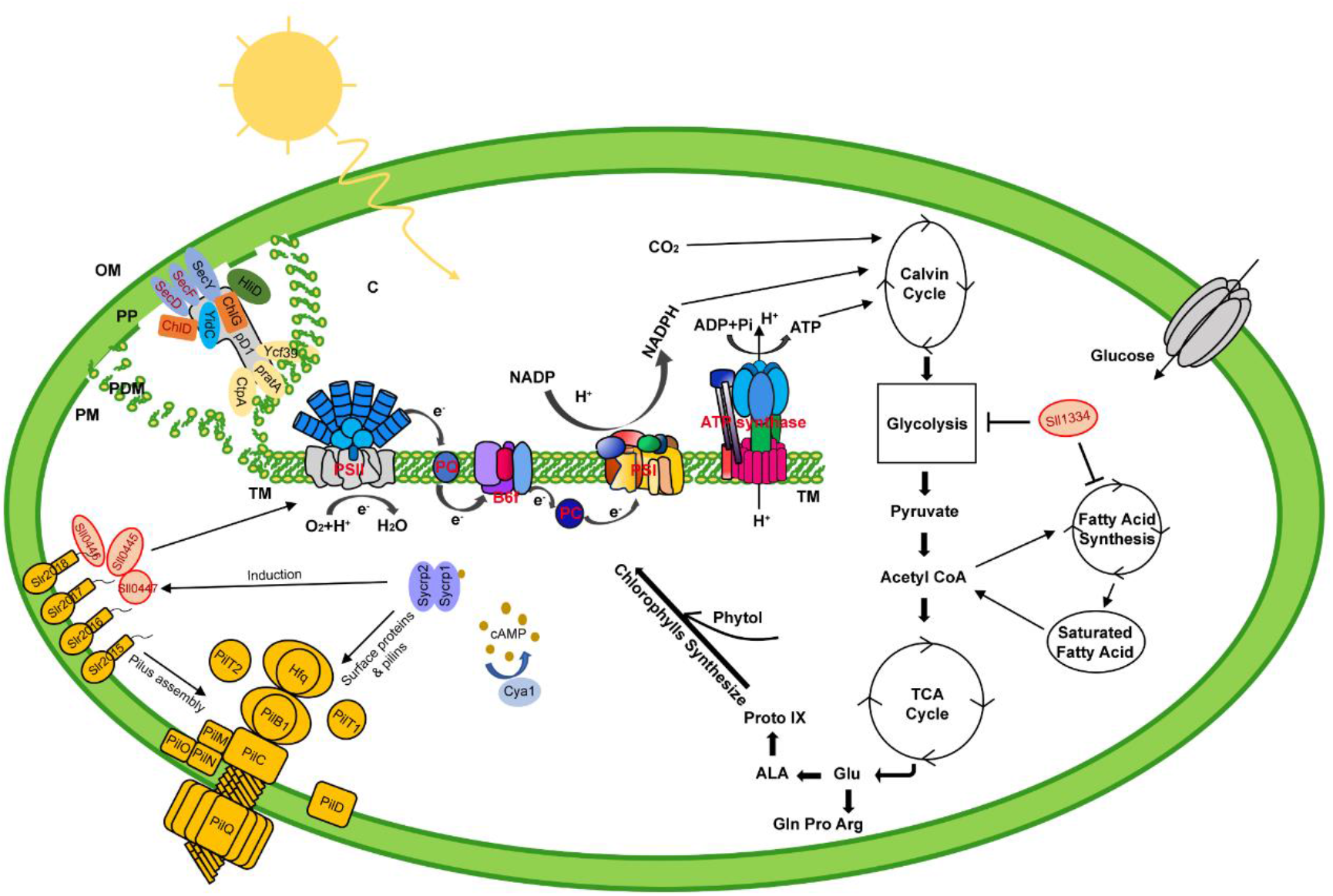
The model of cyanobacterial phototaxis regulation. The model is based on our data (red color) and public knowledge, showing the phototaxis assemble and how it influenced by the structure of pilus and carbon metabolism. Proteins SecD, SecF, and ChlD present a good elution profiling with PSII assembly proteins, indicating that Sec complexes and ChlD might play a critical function in the maturation of PSII. The hypothetical proteins Sll0445, Sll0446, and Sll0447 are induced by cAMP receptor protein Sycrp1 and interact with protein pilus assembly proteins to regulate cell motilities. These hypothetical proteins can also interact with photosynthetic core proteins, revealing a close relationship between photosynthesis and cell motility in *Synechocystis*. The protein Sll1334 can regulate glucolipid metabolism and may have an important role in controlling the utilization of sugars and lipids during heterotrophic growth and then effect the photosynthesis. pD1, precursor of D1; C, cytoplasm; L, lumen; OM, outer membrane; PDM, PratA-defined membrane; PM, plasma membrane; PP, periplasm; TM, thylakoid membrane.

## Conclusion

In this study, combing CoFrac–MS and quantitative proteomics strategies, we are able to predict 291 protein complexes consisting of 24,092 highly confident PPIs in *Synechocystis*, which is the largest protein interaction dataset of *Synechocystis* so far. The comprehensive PPIs information greatly enhanced our understanding of the molecular mechanisms of photosynthesis as well as other fundamental molecular organization in cyanobacterium.

From the predicted PPIs, most of the proteins that tend to have lower degrees involved in metabolic regulation, whereas the proteins with higher degrees are endowed with more basic and conserved functions in *Synechocystis*. By separating photosynthetic proteins interaction networks from whole PPIs, we elucidated how photosynthetic proteins connect with many other functional proteins and influence plenty of necessary biological processes. By comparing protein complexes in *Synechocystis* with other species, including *A.thaliana, E.coli, H.sapiens*, and *S.cerevisiae*, we observed protein complexes evolution and functional variations in different species. For example, the change of NQOs components in different species may be due to that photosynthetic organisms, like *Synechocystis* and *A.thaliana*, had to undertake the process of photosynthesis and recovery from photosynthetic damage. According to the predicted complexes, the hypothetical proteins Sll0445, S0446, and Sll0447 were found to build a functional connection between photosynthesis and cell motility. Photosynthetic apparatus serves as a regulator in energy metabolic process in living cells. Photosynthesis has a close relationship with chemotaxis, because one of the primary purposes of cell motility is to obtain light so that photosynthesis can use light energy to fix CO_2_ for provided energy in the reverse of cell motility. Moreover, the expression of ACC complex was upregulated when the protein Sll1334 was depleted. Cyanobacterium was considered as a promising organism for producing biofuels, but current productivity was still needed further improvement in cyanobacterial systems. The protein Sll1334 was found to be a negative regulator of glycolipid metabolism, which provided us new insights to improve biofuel productivities by genetic modification.

The global landscape of native protein complexes in *Synechocystis* provides a valuable resource for researchers to find and determine the new and promising complexes for further study, and expand our knowledge of protein interaction network, which governs the life of cyanobacteria.

## Materials and methods

### Growth condition and protein extraction

*Synechocystis* strain was grown in liquid BG11 medium at 30 °C in the light (30 μmol m^−2^ s^−1^). The cells were collected by centrifugation (6000g at 4 °C for 5min) when grew to exponential phase (OD730 = 0.8 −1). The lysis buffer contained 20mM Tris-Cl (pH = 7.5), 150mM NaCl, 1%DDM (Catalog No: D4641, Merck, Darmstadt, Germany), and Complete Protease Inhibitors EDTA–free (Catalog No: 4693124001, Roche, Basel, Switzerland) and applied to sonication (5s on, 10s off) for about 5min on ice with an output of 135W. Then the cell debris was discarded by centrifugation (12,000g at 4 °C for 10min). The protein rotein concentration of each sample was measured using the Bradford assay.

### Size–exclusion chromatography, ion-exchange chromatography, and sucrose density gradient centrifugation

*Synechocystis* cell lysate was fractionated by SEC, IEC, and Suc–DGC by using a Thermo Scientific Ultimate 3000 HPLC system. The lysates were injected (350ul per injection) onto MAbPac SEC-1 (Thermo Scientific, 5μm, 300 × 4.0mm) or Superose 6 10/300GL column (GE Life Sciences). There are 24 100ul fractions collected by using MAbPac SEC-1, with flow rate 0.2ml min^−1^. And there are 45 300ul fractions collected by using Superose 6 10/300GL column, with flow rate 0.3ml min^−1^. Protein standards (thyroglobulin, BSA, Albumin egg and myoglobin) were analyzed with the same method to obtain the approximate MW range across fractions. An ion-exchange column (200 × 4.6 mm, 12 μm, 1500 Å, Columnex, San Diego, CA, USA) was also used for lysate samples, and a 110 min salt gradient (0.12M to 1.2 M NaCl) was used to collect 43 fractions. The elution composition %A containing 10mM Tris-HCl (pH = 7.6), 0.5mM DTT and 5% glycerin, while %B with additional 1.2M NaCl. Lysate samples were loaded onto a 12ml 15-70%(w/v) linear sucrose gradient, which was then centrifuged at 160,000×g at 4°C for 16 h in a Beckman MLS–50 rotor (Beckman–Coulter, CA, USA), and 24 fractions were collected. In total, 181 fractions were collected.

### Affinity purification mass spectrometry

The target protein was combination with green fluorescent protein (GFP) on its C– terminal. Cell lysis contained GFP tag strain was subjected to affinity purification by using Anti–GFP antibody (ab290, Abcam, Cambridgeshire, UK). The process of antibody purification was using GenScript Protein A MagBeads according to the manufacturer’s instructions (Catalog No: L00273, GenScript, Piscataway, NJ, USA). Then, the sample was detected by mass spectrometry.

### Trypsin digestion, and peptide clean up

Proteins from all HPLC fractions were precipitated with 10% Trichloroacetic Acid at 4 °C overnight, and dissolved in 50mM ammonium bicarbonate. Trypsin (Catalog No: V5113, Promega, Madison, WI, USA) was added at the ratio of 1:50 and incubated overnight at 37°C. Each fraction desalted using ZipTip C18 plates (Catalog No: ZTC18S960, Millipore, Darmstadt, Germany). Peptides were dried using a LABCONCO evaporate and then resuspended in 0.1% formic acid for further analysis.

### nanoLC–MS/MS analysis

The peptides were dissolved in the 0.1% formic acid, and analyzed using Q–Exactive Plus Orbitrap mass spectrometer (Thermo Fisher Scientific, Waltham, MA, USA). Peptides in 0.1% formic acid were separated on a C18 nano–trap column at a flow rate of 500nL/min. Peptides were ionized at 2.0 kV and using HCD fragmentation of precursor peptides for MS/MS analysis. The MS/MS spectra of the top 20 most intense signals were acquiring by using a data–dependent method. The dynamic exclusion duration was set as 40s and 5 ×10^4^ ions were set to generate MS/MS spectra in the automatic gain control (AGC). The Sequest version 2.1 was used to retrieve the RAW data using a target–decoy based strategy, supplied with the *Synechocystis* 3508 protein database from UniProt database. Up to 2 missed cleavages were allowed.

### Data analysis

R Language and Python scripts were applied to data analysis. Elution profile for individual protein was normalized and smoothed by using scale command in R Language. The Mapp of all proteins identified in our dataset was calculated similar as the previous publication [26]. After that, the ratio of Mapp to the predicted monomeric mass (Mmono) was calculated, and the value of Rapp (Mapp/Mmono) can effectively reflect the oligomerization state of proteins during protein separation process. The protein with a value of Rapp ≥ 2 imply oligomerization state, and Rapp ≤ 0.5 means it may degrade during the protein extraction process and would be discarded in subsequent analysis. The protein with a value of 0.5 < Rapp < 2 means exit as monomeric state.

### Machine learning

We used EPIC software for automated scoring of our data for large–scale determination of high–confidence physical interaction networks and macromolecular assemblies from diverse biological specimens. This software package can be obtained from https://github.com/BaderLab/EPIC. Protein pairs were scored based on five features: MI, Bayes Correlation, Jaccard, Pearson Correlation Coefficient, and Apex Score. We manually collected a data set of “gold standard” protein complexes by the reference database (UniProt) for machine learning analysis, which contains 48 conserved true positive protein complexes. The positive PPIs are defined if they are appeared in the same protein complex, while the components of negative PPIs are from proteins existed in the different protein complex. Then the positive and negative PPIs were used to train the machine learning classifier. The protein pairs that elution profile similarity score more than 0.5 were required and the proteins that used for machine learning were detected with not less than 2 peptide spectrum matches in at least one of the experiments.

### Construction of plasmid

The single mutants of the three *sll0445–sll0447* gene clusters and *sll1334* were generated by inserting a Cm^R^ into their ORFs. The *sll0445* and *slr0149* were cloned with C–terminal GFP–tagged for APMS. The targeted gene and its flanking sequences were amplified by PCR with *Synechocystis* chromosome DNA as the template and cloned into pMD18–T vector (Catalog No: D101A, Takara, Japan). The physical map of the *Synechocystis* genome region containing *sll0445–sll0447* or *sll1334*, indicating the insertion position of Cm^R^ into target gene respectively, is shown in Figure S10B. Primers for mutant construction are as follow:

M_*sll0445up* GTTCAGCGGTGATGAGTVG; M_*sll0445*down GTAAATCAAACAG GGCATG; M_*sll0446up* TGTGGCCTATACAATGTCCCAG; M_*sll0446*down AAGA TATTTCTTCCAGCAAATGG; M_*sll0447*up ATCTCGTATTAAGAAAGCTTG; M_*sll0447*down TGAGCATAAACTGGACTAATG; M_*sll1334up* AGACGGTTAGA ACCAACAGTCACTG; M_*sll1334*down ACAATTTGTAAGCCCTGGCGAACG.

### Cell motility assays and measurement of photosynthetic activity

The phototactic movement was tested according to Wilde et al. [68]. The strains grown on solid BG11 medium containing 0.3% sodium thiosulfate, 8 mM TES (pH = 8.0), 0.8% agar, and 5 mM glucose, under unidirectional illumination with light intensity at 1 −5 μmol photons m^−2^ s^−1^ and the movements were recorded at day 6. The modification of PSII photochemistry in *Synechocystis* was evaluated by OJIP curve, measured by Plant Efficiency Analyzer (Hansatech, Germany).

### RNA isolation and transcriptional analysis

About 50ml of *Synechocystis* grown in BG11 was collected by centrifugation at 4°C and the total RNA was extracted using the TRIzol Reagent (Catalog No: 15596–026, Invitrogen, Waltham, MA, USA). The cDNAs were synthesized with the Perfect Real Time kit (Catalog No: RR047A, Takara, Japan), and used as a template for transcriptional analysis, while the control gene was RNase P subunit B (rnpB). Primers for real–time PCR:

*accA*up AAATGTTTCGGTTAGATGTCC; *accA*down CCAAAGAATAGCCGCACA; *accC*up TTTGGTGGATGGTAACGG; *accC*down TGGCGGAAGCGGAGTTTT; *rnpB*up ACCGCTTGAGGAATTTGGTA; *rnpB*down TTAGTCGTAAGCCGGGTTC T.

## Data availability

All LC/MS/MS Raw data related to this work will be uploaded to iProX (www.iprox.org) and are available with ID IPX0001620001. All other data was uploaded as supplementary material.

## Authors’ contributions

CX performed the experiments with the association of LXY and YW, analyzed the data with the association of BW and LZH, and wrote the manuscript. CX and LY drafted figures. SLC performed Sucrose density gradient centrifugation assays. AE participated in the discussion and guide LZH for data analysis. CHW designed and coordinated the study and contributed to draft the manuscript. All authors read and approved the final manuscript.

## Acknowledgments

We thank Prof. Baosheng Qiu, Prof. Haibo Jiang, and Dr. Guozheng Dai (Central China Normal University) for providing algal, cloning vectors and technical guidance. We would like to particularly thank Prof. Feng Ge (Institute of Hydrobiology, Chinese Academy of Sciences) for the valuable discussion. This work was funded by the Young Thousand Talents Program. We also thank other members of Wan lab for all their support to this project.

