## Supplemental Figure for "Global Landscape of Native Protein Complexes in *Synechocystis* sp. PCC 6803"

### Supplementary material

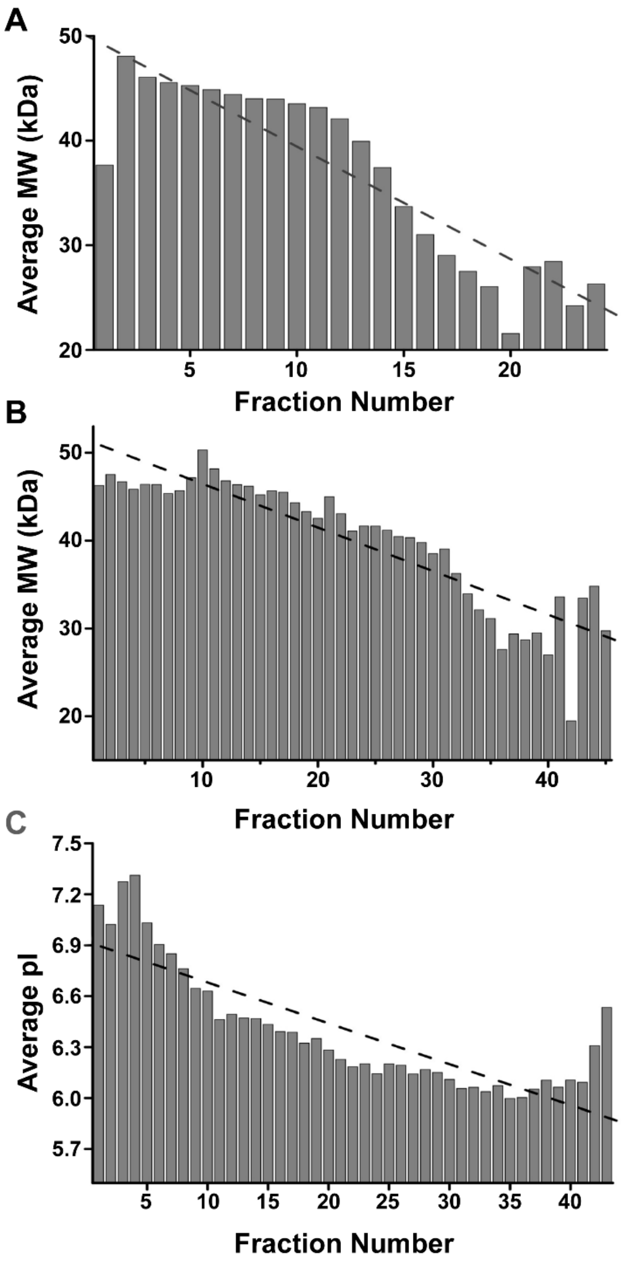


**Figure S1 The distribution of average MW or pI in all fractions**

**A.** MAbPac SEC–1 column. **B.** Superose 6 10/300GL column. **C.** IEC mixed–bed ion–exchange column. For each fraction, the average MW or pI is average MW or pI of all proteins identified in that fraction. The dash lines are trending line.

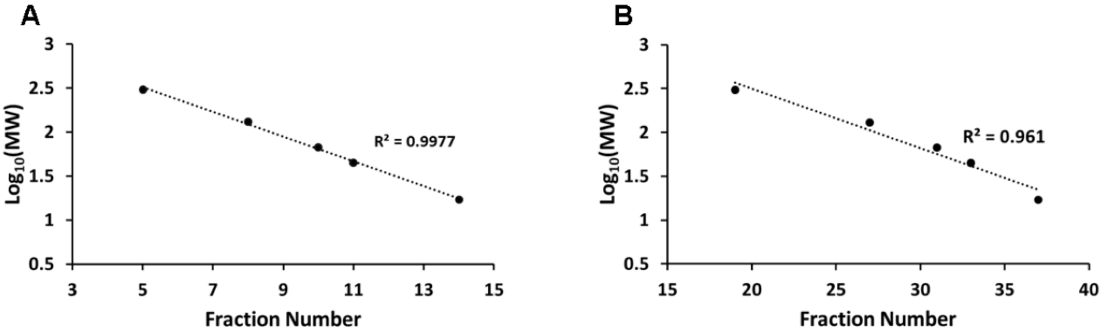


**Figure S2 Regression curve of standard proteins**

Protein standards of known MW (thyroglobulin, BSA, Albumin egg and myoglobin) were separated by SEC column, and their elution peaks were used to calculate approximate MW of the fractions. **A.** MAbPac SEC–1 column and **B.** Superose 6 10/300GL column.

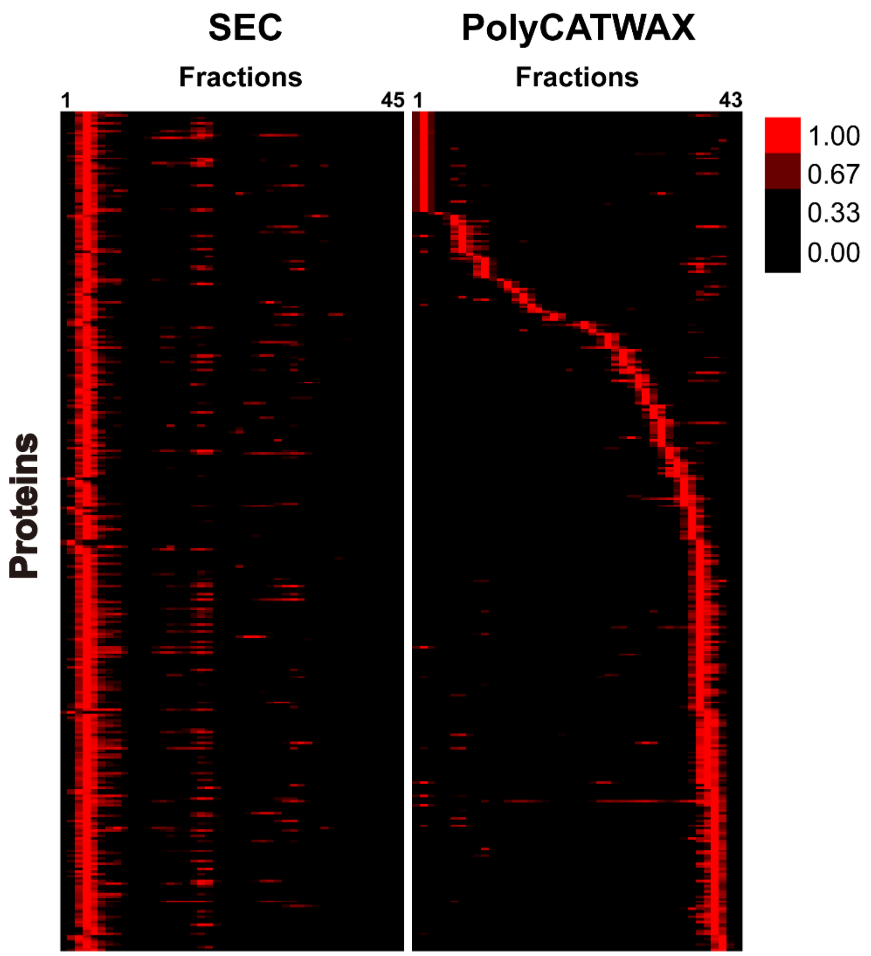


**Figure S3 The complementarity of protein elution profiling in SEC and IEC**

The proteins that not separated effectively in SEC (left) are the protein complexes with MW beyond SEC valid separation range and eluted in early fractions. However, some of these protein complexes can be separated on IEC according to their elution profiling (right). Red color corresponds to protein abundance.

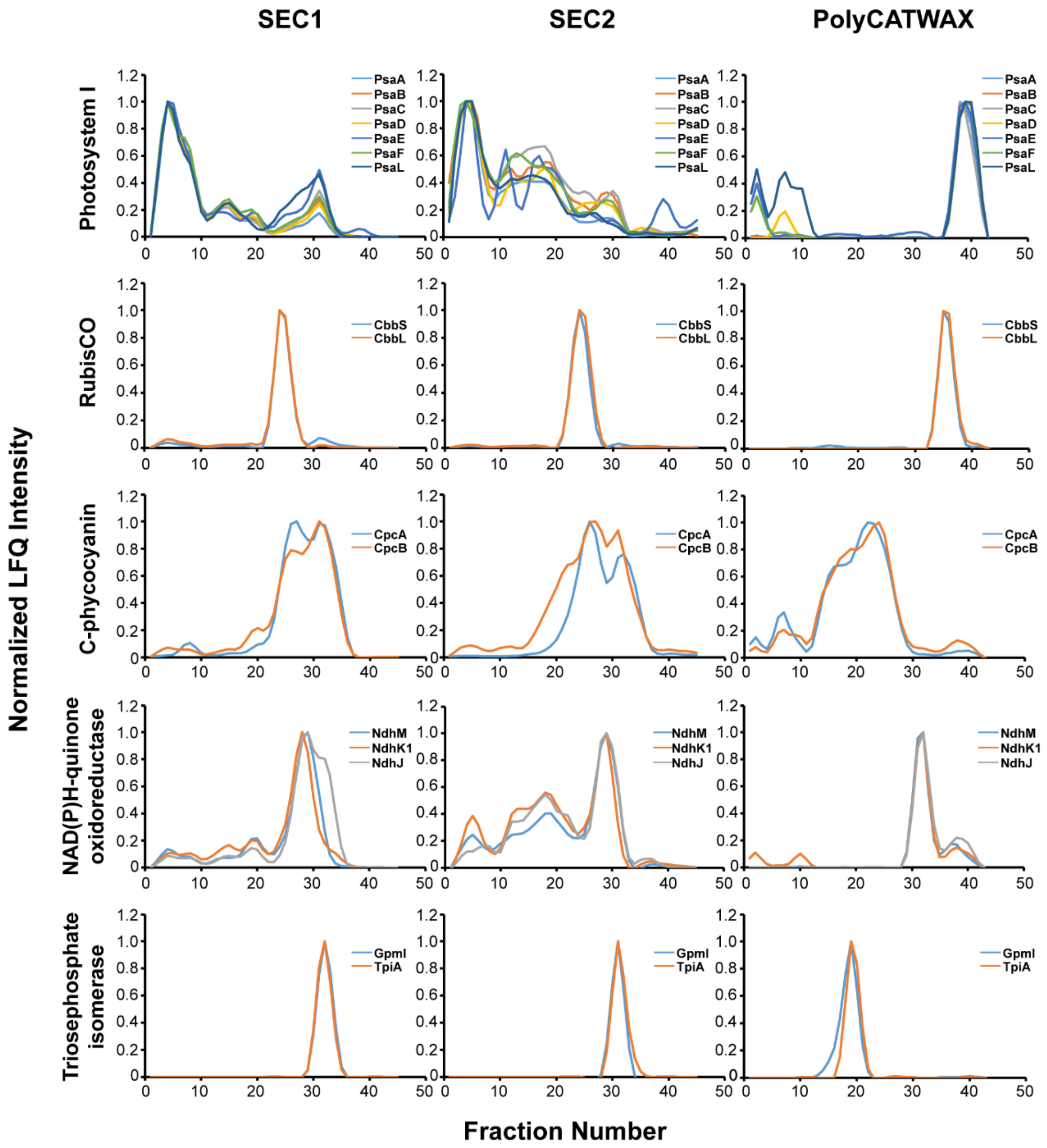


**Figure S4 Elution profiles of components of known protein complexes**

Elution profiling of Photosystem I, RubisCO, C–phycocyanin, NAD(P)H–quinone oxidoreductase and Triosephosphate isomerase on three different columns (SEC1: MAbPac SEC, SEC2: Superose 6 10/300GL, IEC mixed–bed ion exchange). The elution profiling lines of proteins in one protein complex are shown in different colors. x–axis: elution fraction number, y–axis: normalized label–free quantification intensity.

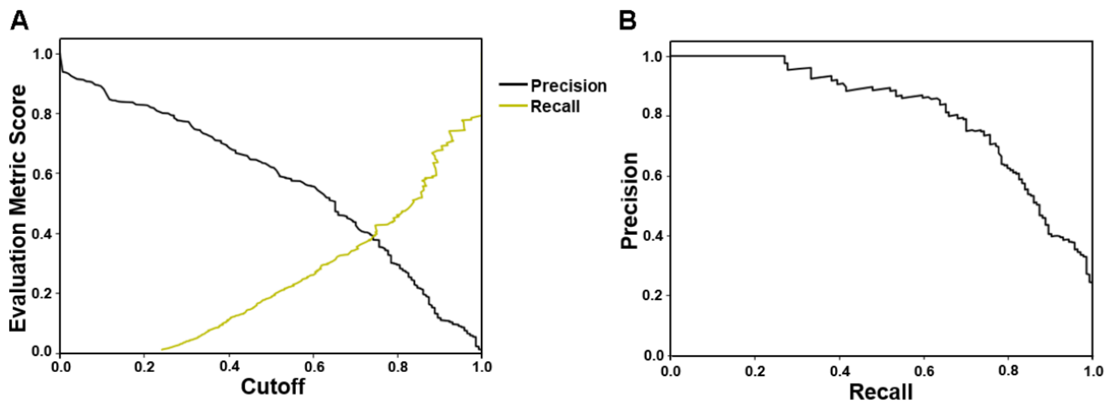


**Figure S5 Evaluation results for machine learning**

**A.** Evaluation metric score–Cutoff from experimental data. **B.** Precision–recall curve (PR) for co–complex PPI prediction from experimental data.

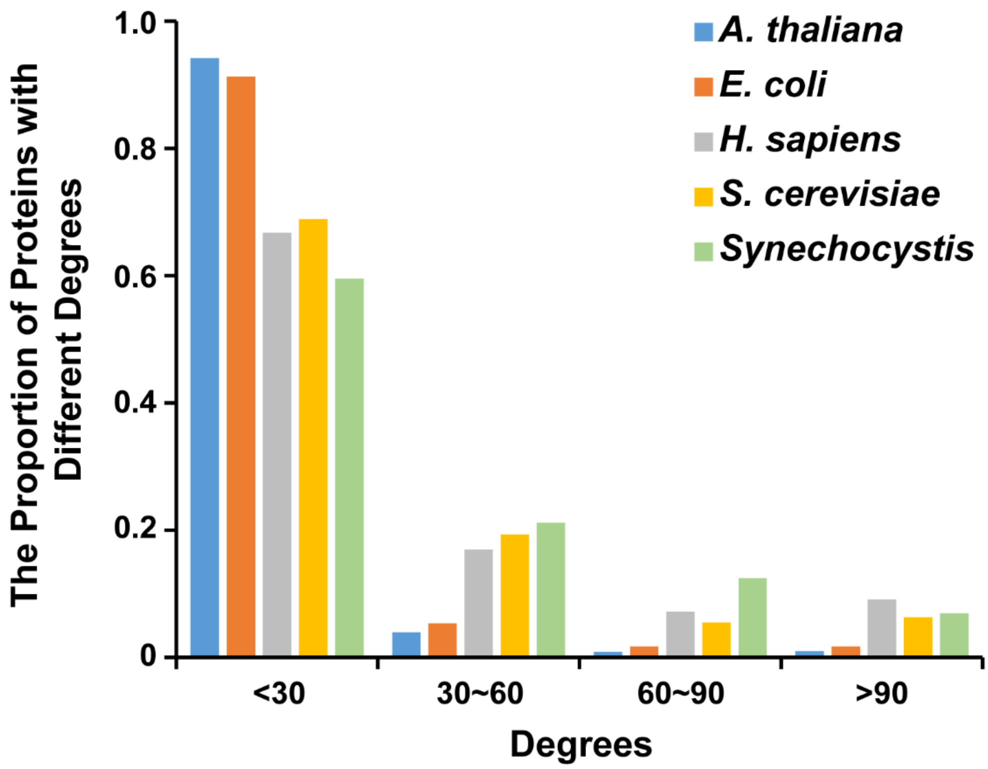


**Figure S6 The distributions of proteins degrees in different organisms**

The protein–protein interaction pairs of *Synechocystis* were generated from our dataset, and PPIs of other model organisms were obtained from the Mentha database. The degree is defined as the number of edges that one protein links to other proteins in the network.

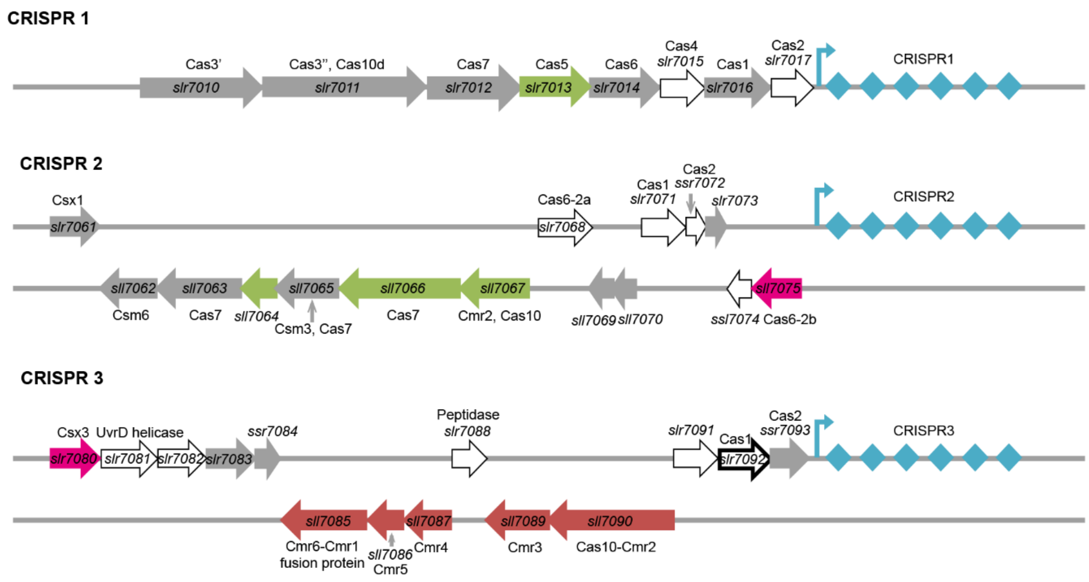


**Figure S7 Organization of the three CRISPR–cas systems in *Synechocystis***

The CRISPR–cas systems were illustrated based on previous work [37,38]. The cas–genes are represented by arrows and located upstream of the CRISPR arrays. Arrows in white represent proteins not identified by MS. Arrows in grey illustrate the proteins, of which no high confident physical interactions were found in our dataset. Other arrows in the same colors represent those proteins that can form complex, such as the CRISPR3 can form one complex (red color), CRISPR2 proteins can have interactions with both CRISPR 1 (green) and CRISPR3 (purple).

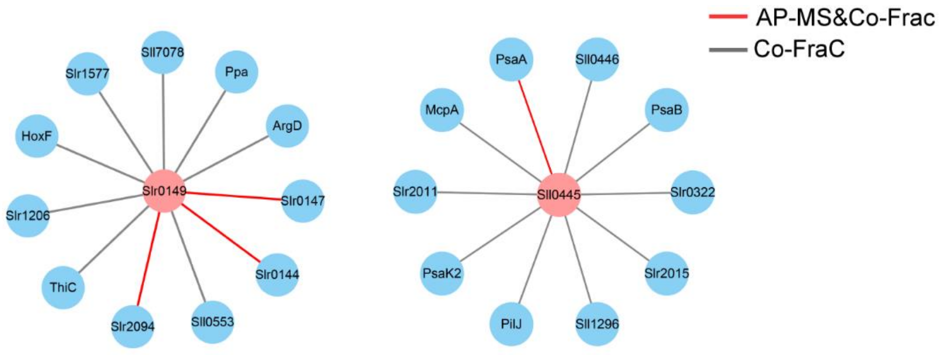


| | Proteins | PSM Count |
|---|---|---|
| **Bait** | **Slr0149-GFP** | **12** |
| Prey | Slr0144 | 6 |
| | Slr0147 | 6 |
| | Slr2094 | 3 |
| **Bait** | **Sll0445-GFP** | **14** |
| Prey | PsaA | 5 |
| | PsaF | 8 |
| | Slr2018 | 8 |

**Figure S8 The APMS result of Slr0149**

Schematic illustration of the PPIs involved with protein Slr0149 from Co–fractionation and APMS data. And the mass spectrometry Peptide–Spectrum Match (PSM) number of the proteins co–purified with Slr0149 from GFP–tagged APMS listed in Table S6.

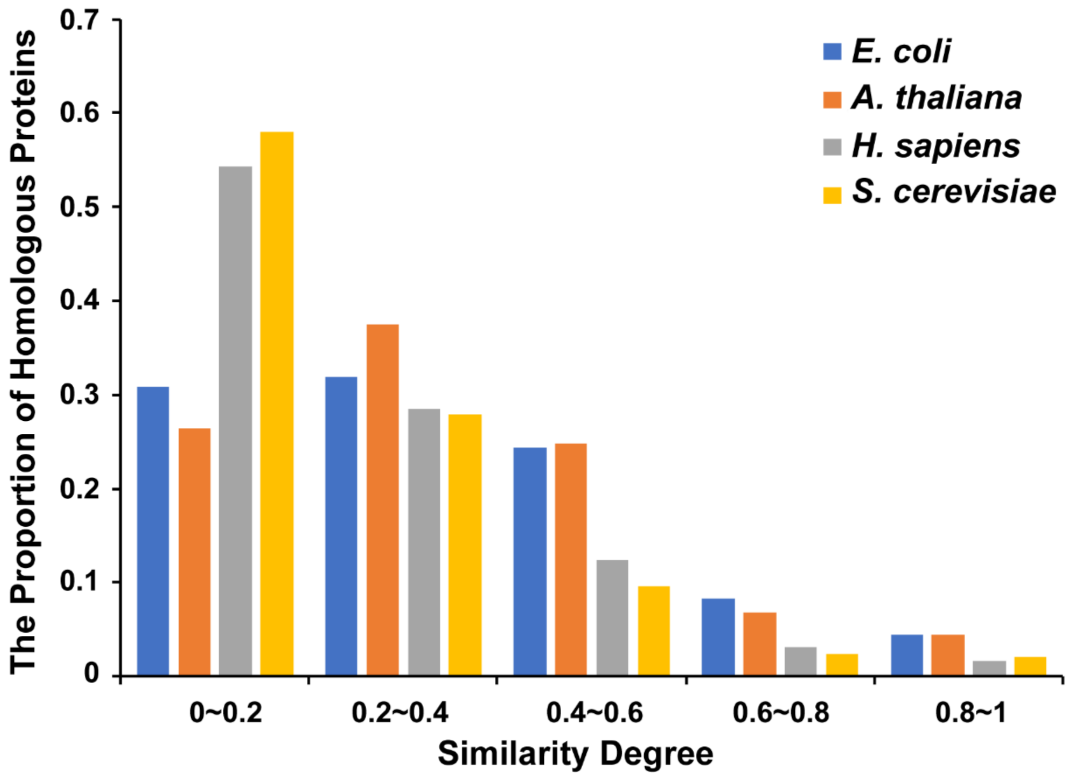


**Figure S9 Conservative analysis of predicted complexes**

The proportion of different similarity degrees of all *Synechocystis* protein complexes in this work was plotted. The similarity degree was calculated by the percentage of protein components in each complex that has homologous in other organisms.

A

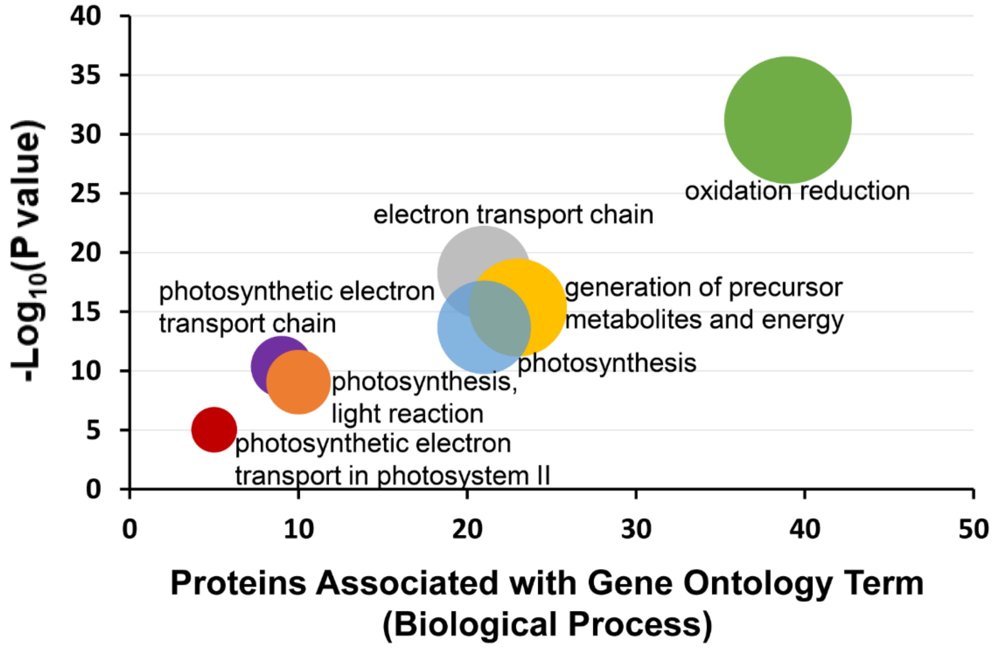


B

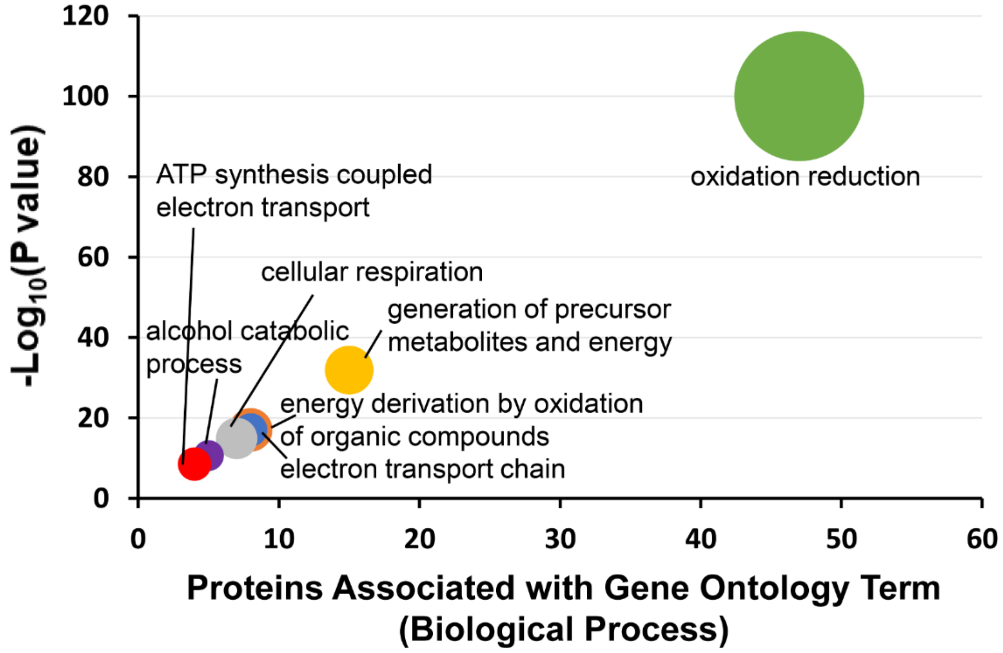


**Figure S10 Comparative analysis of proteins annotated with oxidation–reduction**

Bubble graphs demonstrate the gene ontology terms (biological process) (x–axis) plotted against the $-\log_{10}P$value for oxidation–reduction related protein in homologous with *A.thaliana* part **(A)** and homologous with *A.thaliana* and *E.coli* part **(B)**, respectively.

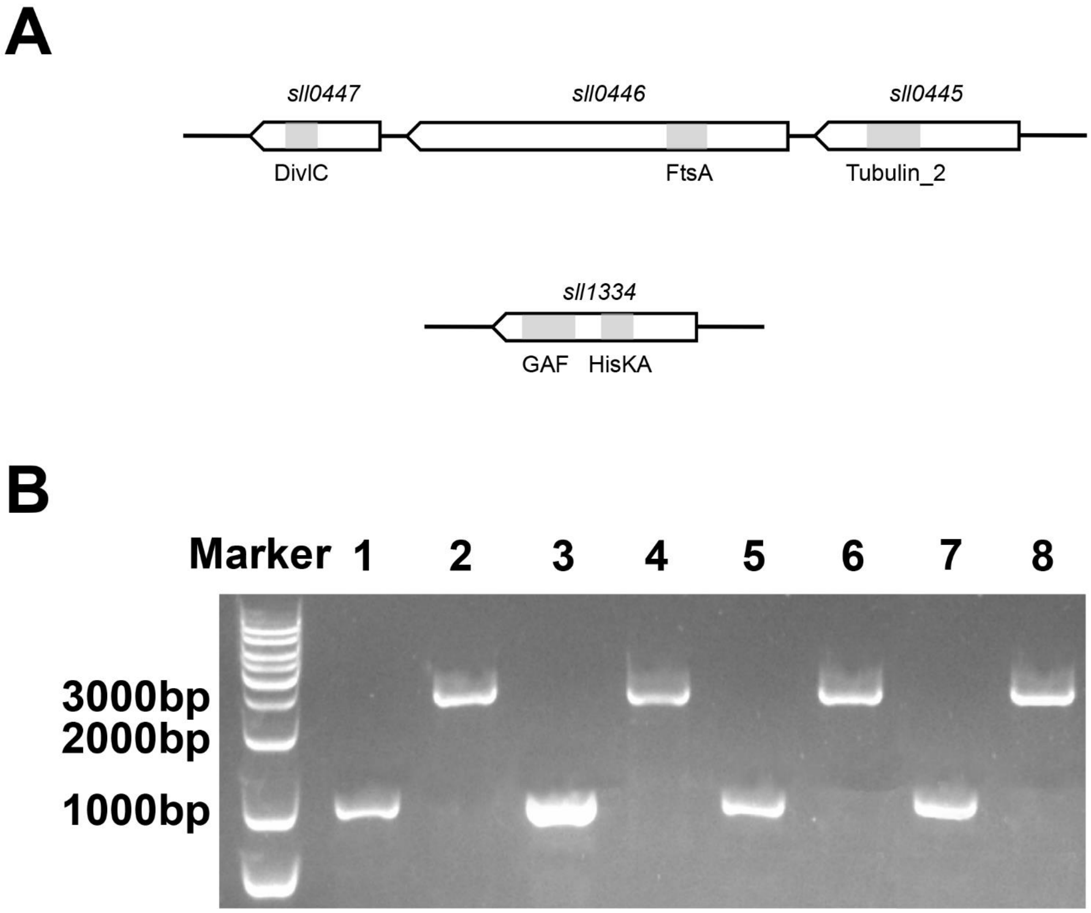


**Figure S11 Construction and detection of the *Synechocystis sll0445–sll0447* gene clusters and s*ll1334***

**A.** The predicted domains of Sll0445–Sll0447 proteins (Top) and Sll1334 (Bottom). The *sll0445* encodes a protein with a Tubulin_2 domain, the *sll0446* encodes a protein with a FtsA domain, and the *sll0447* encodes a protein with a DivIC domain. The *sll1334* encodes a protein with GAF and HisKA domains. **B.** Detection of the degree of segregation of the *sll0445–sll0447* mutants by PCR amplification. Lanes 1–2 use primers *sll0445* up and *sll0445* down. Lanes 3–4 use primers *sll0446* up and *sll0446* down. Lanes 5–6 use primers *sll0447* up and *sll0447* down. Lanes 7–8 use primers *sll1334* up and *sll1334* down. Lanes 1, 3, 5, and 7: wild type strain DNA; lane 2: *sll0445*::$Cm^R$ mutant DNA; lane 4: *sll0446*::$Cm^R$ mutant DNA; lane 6: *sll0447*::$Cm^R$ mutant DNA; lane 8: *sll1334*::$Cm^R$ mutant DNA.
